# BMP-treated human embryonic stem cells transcriptionally resemble amnion cells in the monkey embryo

**DOI:** 10.1101/2021.01.21.427650

**Authors:** Sapna Chhabra, Aryeh Warmflash

## Abstract

Human embryonic stem cells (hESCs) possess an immense potential to generate clinically relevant cell types and unveil mechanisms underlying early human development. However, using hESCs for discovery or translation requires accurately identifying differentiated cell types through comparison with their in vivo counterparts. Here, we set out to determine the identity of much debated BMP-treated hESCs by comparing their transcriptome to the recently published single cell transcriptomes of early human embryos in the study Xiang et al 2019. Our analyses reveal several discrepancies in the published human embryo dataset, including misclassification of putative amnion, intermediate and inner cell mass cells. These misclassifications primarily resulted from similarities in pseudogene expression, highlighting the need to carefully consider gene lists when making comparisons between cell types. In the absence of a relevant human dataset, we utilized the recently published single cell transcriptome of the early post implantation monkey embryo to discern the identity of BMP-treated hESCs. Our results suggest that BMP-treated hESCs are transcriptionally more similar to amnion cells than trophectoderm cells in the monkey embryo. Together with prior studies, this result indicates that hESCs possess a unique ability to form mature trophectoderm subtypes via an amnion-like transcriptional state.

## Introduction

Human embryonic stem cells (hESCs) provide a unique window into early stages of human development. Over the last few years, they have been used to generate many medically relevant cell types and models of early human development [1,2]. However, lacking the spatial context that the embryo provides, the in vivo identity of cells obtained from differentiating hESCs is often unclear. The identity of BMP treated hESCs has been particularly controversial, with arguments made for three different extra-embryonic tissues - trophectoderm, amnion and extra-embryonic mesoderm [3–5]. Based on limited transcriptional data from the human and monkey embryo, we previously argued that BMP-treated hESCs are more likely to represent trophectoderm cells than extra-embryonic mesoderm cells ([6]). However, it was not possible to make a direct comparison with human amnion cells due to the lack of in vivo data.

Obtaining data directly from human embryos is of paramount importance because there are significant differences between human embryos and those of mammalian model organisms such as the mouse, especially in the formation of amnion - the extra-embryonic tissue that covers the embryo in a protective sac [7,8]. In human and monkey embryo, the amnion is formed prior to gastrulation, whereas in mouse it is formed after gastrulation and is partially derived from primitive streak cells [8,9]. There have been no reports on the molecular characterization or lineage relationships of the amnion in humans until recently.

In a major breakthrough, a recent study (Xiang et al 2019) succeeded in obtaining the transcriptional signature of cultured human embryos in the second week of embryonic development [10]. This study provided transcriptomes for all major cell types in the human embryo from embryonic day 6 to 14 (D6-D14) and included the first transcriptomes of putative amnion cells (2 cells at D12 and 11 cells at D14).

To discern the in vivo identity of BMP-treated hESCs, we first reexamined whether the data in Xiang et al. support labeling the cells denoted as amnion as a distinct cell type as prior studies have hinted on the transcriptional similarity between amnion and trophectoderm cells. Monkey amniotic cells in vivo or purported human amnion cells in vitro express TFAP2A, GATA2/3, CDX2, and TP63, all well-known trophectoderm markers [4,11,12]. Surprisingly, Xiang et al neither examined the transcriptional similarity of the two fates nor provided a rationale for assignment of amnion fate to cells.

Our analyses revealed that cells labelled as amnion comprise a mix of different cell types, most of which are indistinguishable from syncytiotrophoblast cells. The mislabeling in the Xiang et al study can be attributed to the inclusion of pseudogenes in that analyses. In the absence of a molecular signature for the human amnion, we turned to the recently published monkey embryo transcriptome [13] to resolve the identity of BMP-treated hESCs. Comparing the transcriptional signature of BMP-treated hESCs with early post-implantation monkey amnion and trophectoderm cells revealed that they are more similar to monkey amnion cells. Together with prior studies that have revealed the functional similarity of BMP-treated hESCs with human trophectoderm cells ([3,14]), this result potentially hints at an ability of hESCs to differentiate into trophectoderm cells through an intermediate amnion-like transcriptional state. Our analyses also revealed additional mislabeled cellular populations in the Xiang et al dataset. Notably, the cells identified as a novel intermediate cell type likely represent extra-embryonic mesodermal cells, a transient extra-embryonic cell population that also develops prior to gastrulation in the human and monkey embryo [9,15,16]. Additionally, putative inner cell mass cells are likely mislabeled cytotrophoblast cells. In summary, our analysis reveals the transcriptional similarity of BMP-treated hESCs with early post implantation monkey amnion, provides a corrected dataset based on the work of Xiang et al. that can be used to study early human development, and suggests that more work will be needed to identify the in vivo transcriptome of human amnion.

## Results

### Putative amnion cells express trophoblast specific lineage genes

Although primate amnion is presumably derived from epiblast cells [8], both monkey amnion cells and hESC derived putative amnion cells exhibit transcriptional similarity with trophectoderm cells [4,11,12]. To determine the similarity of amnion with epiblast and trophectoderm lineages, we compared the expression of lineage specific genes between individual putative amnion cells in the Xiang et al dataset with the average expression of these genes in cells corresponding to the three lineages – the epiblast, primitive endoderm and the trophectoderm.

We utilized the lineage-specific genes documented in the Stirparo et al 2018 study [17], which consolidated data from previous studies [18–24] and identified a group of 12 high confidence lineage specific genes – NANOG, SOX2, KLF17, TDGF1, PDGFRA, GATA6, GATA4, SOX17, GATA3, GATA2, KRT18, TEAD3 that effectively separate the three lineages of the pre-implantation human embryo [17]. We replaced KRT18 with another well-known trophectoderm marker KRT7 [21], as the latter was more specific to trophectoderm lineages in pre and peri-implantation stage embryos in Xiang et al 2019 dataset (Fig S1D).

This known lineage marker gene set effectively separates the three lineages, even at the post-implantation stage, in both principal component and correlation analyses (Fig S1 C, E, F). It also correctly placed the derived cell types with their respective parent lineages – syncytiotrophoblast (STB) and extra-villous cytotrophoblast (EVT) cells with cytotrophoblast cells (CTB) and primitive streak cells with epiblast cells.

In tSNE analyses presented in Xiang et al 2019, D12 amnion cells are placed with epiblast cells, while the D14 amnion cells are placed in the PSA (primitive streak anlage) cluster, indicating transcriptional similarity of amnion with epiblast and primitive streak cells (Xiang et al 2019 Figure 2; Fig 1A). However, our analyses with lineage specific genes contradicts this result and instead shows transcriptional similarity of D14 amnion cells with trophectoderm cells, not with epiblast or primitive streak cells (Fig S1E). Consistent with this, most D14 amnion cells do not express known pluripotency and primitive streak markers, thus questioning their placement in the PSA cluster (Fig 1B, C).

**Fig 1:**
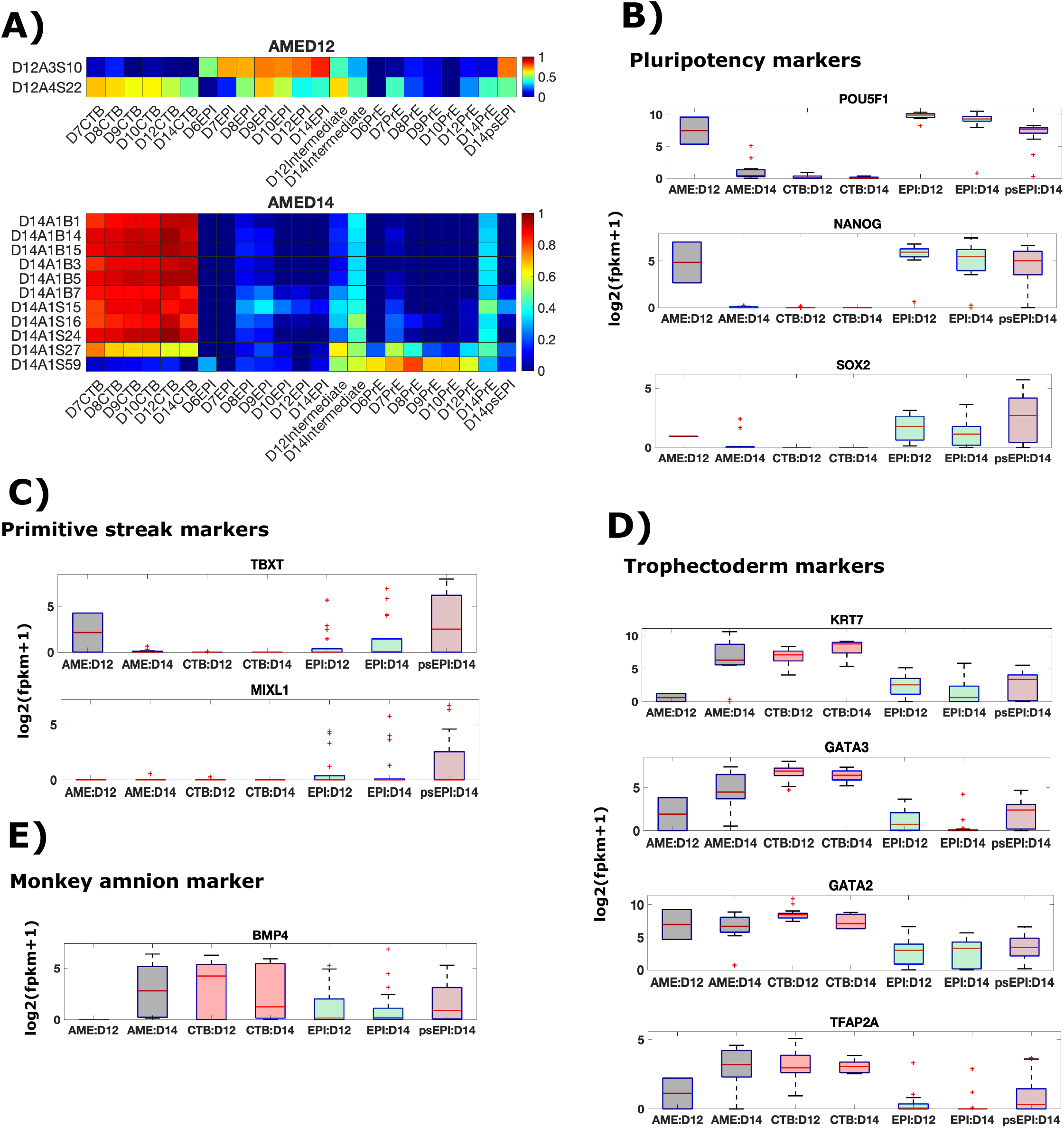
Putative human amnion cells express trophoblast lineage specific genes. ***(A)*** Heatmap showing Pearson correlation coefficients for expression of known lineage markers between individual amnion cells and indicated cell type averages. ***(B, C, D)*** Box plots showing expression of indicated genes in indicated lineages.

**Fig 2:**
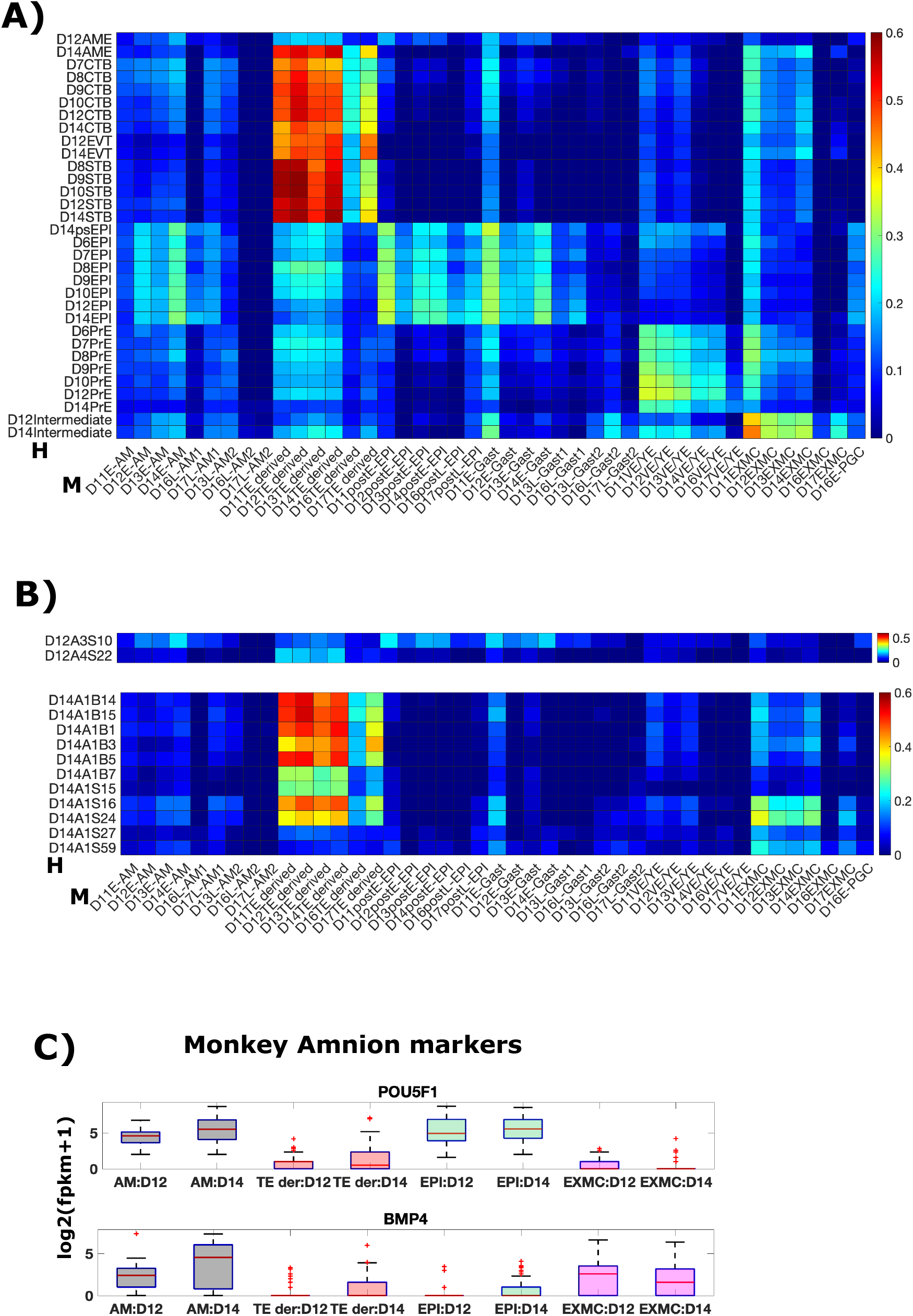
Putative human amnion cells are transcriptionally more similar to monkey trophectoderm-derived than monkey amnion cells. ***(A)*** Heatmap showing Pearson correlation coefficients of average expression of variable genes in the monkey embryo ((C, E); CV>1 across 1453 monkey cells; 2440 genes) in indicated cell types. ***(B)*** Heatmap showing Pearson correlation coefficients for expression of variable genes in the monkey embryo between individual human amnion cells and monkey cell type averages. The symbols H and M represent the parent embryo of cells - human(H), monkey(M). Genes used I (A) and (B) are the same as those in Fig S2D. ***(C)*** Box plots showing expression of indicated genes in indicated lineages.

Most amnion cells (11/13; 1/2 D12, 10/11 D14) are transcriptionally correlated with CTB cells (Fig 1A). Consistent with this, most D14 amnion cells express trophectoderm markers – KRT7, GATA2/3, TFAP2A at levels comparable to D14 CTB cells (Fig 1D). Strikingly, although the putative amnion cells express high levels of KRT7 in the scRNA seq data, only the trophectoderm but not the amnion, was positive for KRT7 in immunostaining in the same study (Xiang et. al 2019 Figure 1J CK7/KRT7 stain). This suggests that the cells labelled as amnion either post-transcriptionally repress KRT7 or represent mislabeled CTB/CTB-derived cells.

### Putative amnion cells transcriptionally more similar to monkey trophectoderm-derived cells than monkey amnion cells

To further discern the identity of putative human amnion cells, we compared the transcriptomes of cells in the human embryo in the Xiang et al study with cells in the post-implantation cynomolgus monkey embryo [13], which has a very similar morphology as that of the human embryo in the peri-implantation stages [15]. Remarkably, known lineage markers in human embryo also delineate the three lineages in the monkey embryo, highlighting conserved expression of these genes across the two species (Fig S2A, B, C).

In the known lineage gene space, most monkey cell types exhibit transcriptional similarity with their parent or sibling lineages. Amnion and gastrulating cells (primitive streak cells) are transcriptionally similar to epiblast cells, which is their parent lineage (Fig S2C). Most amnion cells, however, are also transcriptionally similar to trophectoderm-derived cells, consistent with the expression of trophectoderm-specific genes in the monkey amnion [11]. Extra-embryonic mesodermal cells (EXMC), which overlay the amnion and develop prior to primitive streak formation in primates [15,16], are transcriptionally similar to visceral/yolk sac endoderm (VE/YE), consistent with them being primitive endoderm derivatives [25].

As this restricted lineage gene space does not distinguish the amnion and trophectoderm lineages, we compared the expression of genes variable across all monkey cells (CV > 1, 1453 cells; 2440 genes). In this space, the amnion cells retain transcriptional similarity with the epiblast but lose similarity with the trophectoderm (Fig S2D). Thus, expression of these genes can be utilized to determine whether putative human amnion cells represent epiblast derived amnion cells, as suggested by Xiang et al 2019, or mislabeled trophectoderm cells, as suggested in the previous section (Fig 1A-D, S1E, F).

Comparing the expression of genes with variable expression in the monkey embryo with mean expression of same genes in the human embryo reveals that the putative human amnion cells are transcriptionally most correlated with monkey trophectoderm-derived cells, not with monkey amnion, further challenging the identities assigned to these cells in Xiang et al 2019 (Fig 2A). This correlation is retained at the level of single human amnion cells, where most amnion cells (10/13; 1/2 D12, 9/11 D14) show highest correlation with monkey trophectoderm cells (Fig 2B). Consistent with this, human amnion cells express trophectoderm genes as shown previously (Fig 1D).

Notably, putative human amnion cells do not exhibit the BMP4+/POU5F1+ (OCT4+) transcriptional signature of the monkey amnion (Fig 2C [11,13]). D12 human amnion cells do not express BMP4. D14 amnion cells express BMP4 comparable to D14 CTB cells and POU5F1less than D14 epiblast cells (Fig 1E, A). This is contrary to their monkey counterparts which express BMP4 higher than trophectoderm-derived cells and POU5F1 comparable to epiblast cells (Fig 2C).

To sum, the transcriptional similarity of putative human amnion with monkey trophectoderm-derived cells and not monkey amnion cells, supports the notion that they represent mislabeled trophectoderm cells.

### Pseudogenes leads to the misclassification of putative amnion cells

We next sought to understand the reason that the analyses in Xiang et al mistakenly classified cells as amnion, rather than trophectoderm. To delineate lineages in the human embryo in a larger gene space, we performed a principal component analyses (PCA) using expressed genes (FPKM>1 in at least 50% of cells within a lineage assigned in the Xiang et al 2019 study) with high variability (CV > 0.5) across all 555 cells. Color coding cells with the lineages assigned in the Xiang et al study reveals that the first principal component separates trophectoderm, primitive endoderm and epiblast cell types while the second principal component separates the putative amnion, intermediate and primitive streak cells from the rest. Restricting the PCA to more variable genes (CV>1, CV>1.5) puts most of the amnion, intermediate and primitive streak cells together on PC1, distinct from the rest of the cells (Fig 3A). This clustering result is broadly similar to the one shown in Figure 2B of Xiang et. al 2019, where the PSA cluster in Figure 2B places D14 amnion, intermediate and primitive streak cells together, distinct from the rest of the cells.

**Fig 3:**
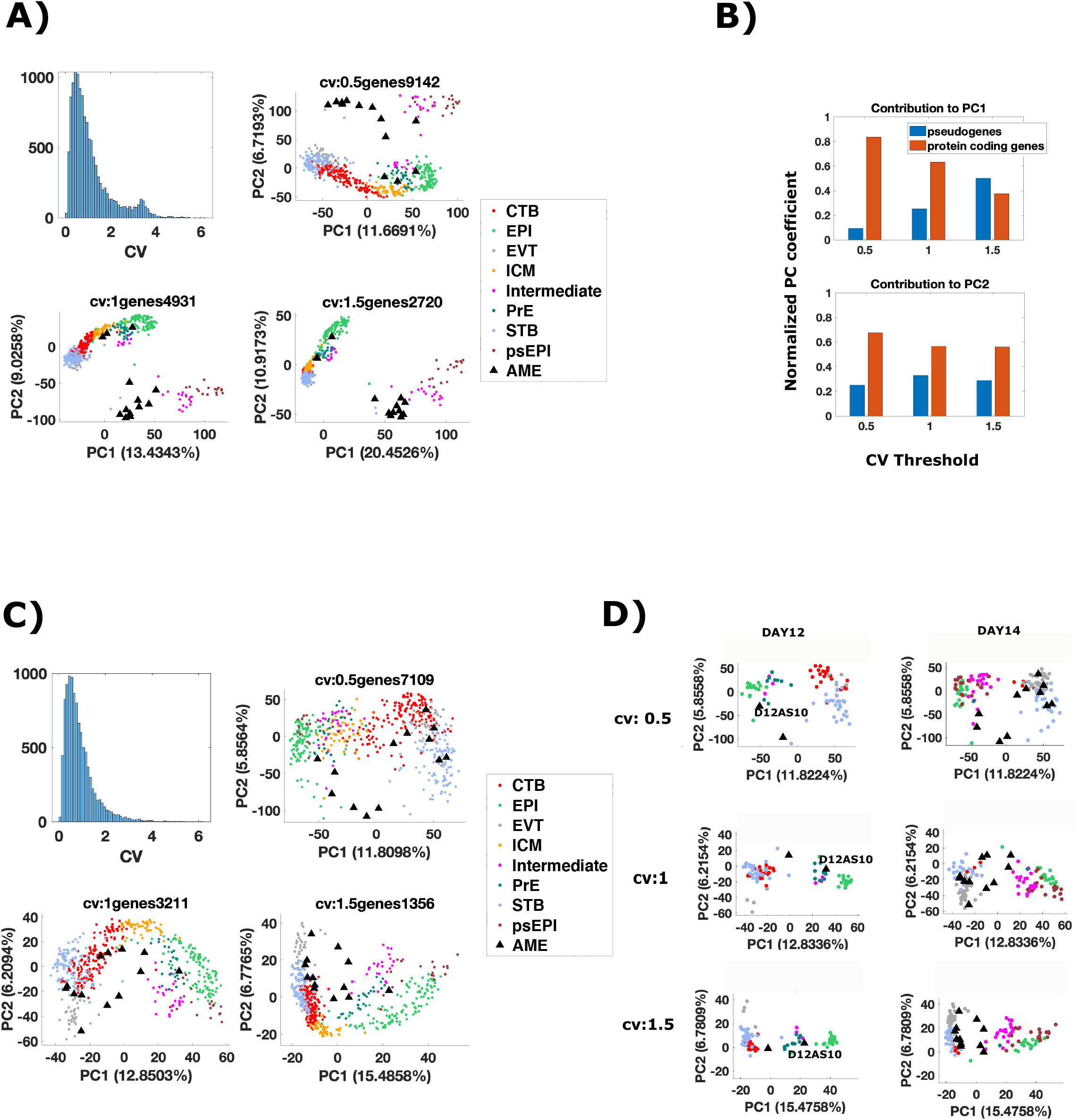
Pseudogenes lead to placement of putative human amnion cells in the PSA cluster. ***(A, C)*** Histogram of Coefficient of Variation of expressed genes (Genes with FPKM>1 in at least 50% cells of a given lineage) across all cells. Principal component analyses (PCA) of 555 cells with variable expressed genes. Variability is defined by a threshold in CV values. The threshold CV and the number of genes that cross the threshold are indicated above the graph. The percentage in x/y labels represents the % of variance in the data explained by each PC. Read counts calculated as log2(fpkm+1) were used for the PCA. Cells are color-coded by the lineages assigned in Xiang et al 2019. In (C), the genes were filtered to include only protein coding genes, prior to determining variable expressed genes. ***(B)*** Contribution of different gene biotypes to the first two principal components. Normalized PC coefficient = Cumulative PC coefficient for a biotype/ Cumulative PC coefficient for all biotypes. Cumulative PC coefficient is calculated as the sum of the absolute PC1 coefficient of all genes in that biotype. The two biotypes that contribute the most are shown. ***(D)*** PCA plots in (C) plotted for a subset of cells corresponding to embryonic day 12 and 14.

To determine the gene categories (Ensembl biotypes) that contribute the most to the two principal components in the above analyses, we plotted the normalized PC coefficient of different gene categories for each principal component. The top two contributors are protein coding genes and pseudogenes. Strikingly, the contribution of pseudogenes increases as the amnion-intermediate-primitive streak cluster moves to a distinct PC1 (Fig 3B).

Pseudogenes are homologous to protein coding genes but with a frameshift or stop codon, which renders them non-translational [26]. While there is some speculation on the role of pseudogenes in gene regulation [27], there is no conclusive evidence for an essential role of pseudogenes in early mammalian development. Hence, we repeated the PCA with only protein coding genes under the same gene selection criteria as before. Contrary to the previous analyses, the amnion cells are now distinct from the intermediate and primitive streak cells (Fig 3C). Instead, most of the amnion cells (10/13 in CV>0.5, >1.0; 7/13 in CV>1.5) share PC1 with trophoblast cells. Plotting the D12 and D14 data separately shows that even with the most restricted gene set (CV>1.5), most (7/11) D14 amnion cells share PC1 with trophoblast cells (Fig 3D). This indicates that transcriptional similarity of protein coding genes is very high between putative amnion and trophoblast cells, consistent with previous section, and their placement with primitive streak cells in the analyses of Xiang et al is due to similar expression of pseudogenes.

### Putative amnion cells contain a mix of EVT, STB and ambiguous cells

To further determine which trophoblast cell type the cells mislabeled as amnion correspond to, we repeated the principal component analyses using expressed genes with high variability (CV>0.5) between D12,14 amnion, CTB, STB and EVT cells (Fig 4A). We removed one putative amnion cell (D12AS10) from this analysis, as it exhibits high correlation of lineage specific gene expression with epiblast cells and is placed either in the primitive endoderm or epiblast cluster in all PCA plots (Fig 1A, 3D).

**Fig 4:**
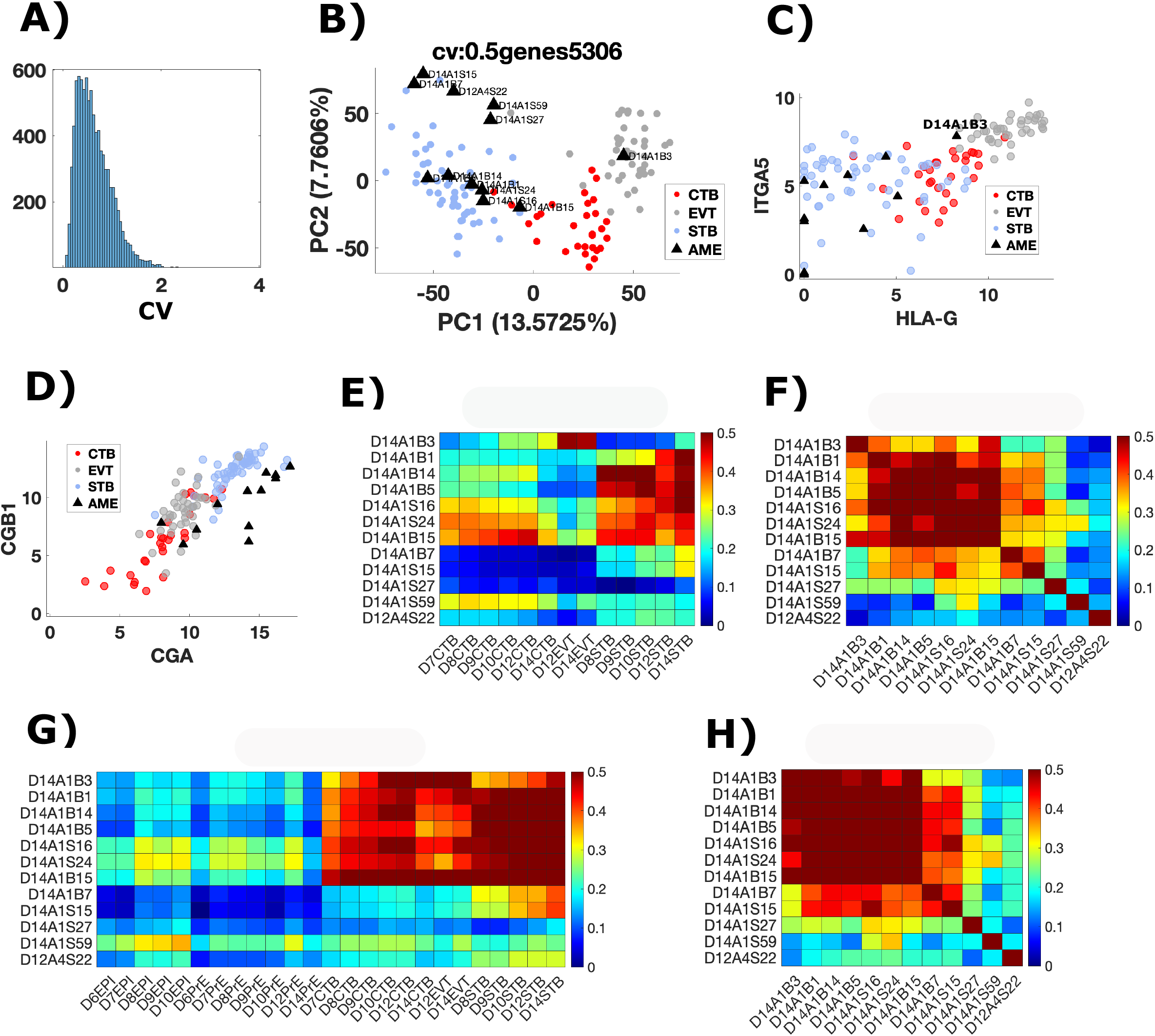
Putative human amnion cells contain a mix of EVT, STB and ambiguous cells. ***(A)*** Histogram of coefficient of variation of expressed genes in D12, D14 CTB, EVT, STB and amnion cells. ***(B)*** PCA of variable genes (CV>0.5) across cells in D12, D14 CTB, EVT, STB and amnion cell types. Methodology and axis labels same as in Fig3 ***(C, D)*** Scatter plots showing expression of indicated genes (log2 (fpkm+1)) in D12,14 CTB, EVT, STB and amnion cells. ***(E, G)*** Heatmap showing Pearson correlation coefficients for expression of variable genes (E: genes used for PCA in Fig 4B; G: genes used for PCA in Fig3C (CV 0.5)) between individual amnion cells and indicated cell type averages. ***(F, H)*** Heatmap showing Pearson correlation coefficients for expression of variable genes (F: genes used for PCA in Fig 4B; H: genes used for PCA in Fig3C (CV 0.5)) between individual amnion cells.

PCA reveals three distinct populations of putative amnion cells (Fig 4B). One cell is placed with EVT cells whereas the others are divided into two groups, both of which comprise STB cells. These two groups might represent different stages within STB maturation. The putative amnion cell in the EVT cluster expresses known EVT markers – HLA-G and ITGA5 [12,28], whereas other putative amnion cells do not (Fig 4C). All putative amnion cells express high levels of human chorionic gonadotrophins genes (hCGA, hCGB1), comparable to STB cells (Fig 4D, [12,28]). Based on hCG protein staining in Extended Data Figure 4u of Xiang et. al 2019, they argue that amnion cells express hCG protein, however, the data shown in that figure is unclear. The cells that have high hCG lie outside a layer of cells surrounding the amniotic cavity, and likely represent STB cells. To distinguish between the two cell types, it is necessary to show an overlap with other known markers. Moreover, hCG is a secreted protein, so its presence in a cell need not imply production in the same cell. Thus, hCG immunostaining alone is not a good indication that it is expressed by amnion cells.

Consistent with PCA, a correlation analyses of expression levels in same gene set also reveals three distinct population of putative amnion cells. One cell (1/12) is transcriptionally correlated (Pearson correlation coefficient > 0.4) with EVT cells, half of the putative amnion cells (6/12) are correlated with STB cells, and the rest (5/12) show either low or no correlation with any of three trophoblast cell types (Fig 4E). To determine if this third population (5/12) comprises a distinct cell type, we examined the pairwise correlation for gene expression of the same gene set within individual amnion cells (Fig 4F). The heterogeneity within these five cells, highlighted by low cell-to-cell correlation values, argues against this. Repeating the correlation analyses for a larger gene set (CV>0.5 across all cells), reveals that most putative amnion cells (9/12) are transcriptionally correlated with one or more trophoblast lineages, but three cells (D14A1S27, D14A1S59, D12A4S22) still remain transcriptionally distant (Fig 4G, H). Within the known lineage marker gene space, D14A1S27, D14A1S59, D12A4S22 correlate with CTB/EPI, CTB and PrE cells respectively but in a larger gene space the lineage relationship is lost (Fig 1A, 4EH). Due to this apparent contradiction, we cannot conclusively determine an identity for these three cells. Amongst the rest, we classify 8/9 cells as mislabeled STB and 1 cell as mislabeled EVT cell. Taken together, our results suggest that the data in Xiang et al 2019 are not sufficient to determine the transcriptome of human amnion.

### BMP-treated hESCs are transcriptionally more similar to early post-implantation monkey amnion than monkey trophectoderm

In the absence of a unique human amnion transcriptome, we turned to the post implantation monkey embryo to resolve the identity of BMP-treated hESCs. We have previously shown that sparsely seeded hESCs treated with BMP4 ligands for 42h transcriptionally resemble trophectoderm cells, and not extra-embryonic mesoderm cells [6]. In this section, we revisited that data and compared the transcriptional similarity of BMP-treated hESCs with monkey amnion and trophectoderm lineages.

We first defined a set of lineage specific genes for the early post implantation monkey amnion, trophectoderm and epiblast (D11-14), and then compared the expression of those genes in monkey embryo with BMP-treated hESCs. We chose early stages (D11-14) of monkey post implantation development because the transcriptional similarity with corresponding human stages is higher at these stages compared to the later (D16-17) (Fig 2A).

To determine lineage-specific genes, we extracted genes that are differentially expressed between that lineage and at least one of the other 2 lineages (fold change > 5, false discovery rate [FDR] = 0.01). From this list, we excluded genes that are differentially expressed between different time points within that lineage (embryonic day [D]11-14) to reduce noise within the lineage and further removed genes with a low expression value (Fragments Per Kilobase of transcript per Million mapped reads [FPKM] < 5 in at least 2 of the 4 time points for that lineage). This gave a list of 571 lineage-specific genes (S1 Table). These genes clearly separate the three lineages transcriptionally (Fig 5A). Examining the genes differentially upregulated in the amnion and trophectoderm compared to the epiblast reveals that the monkey amnion differentially upregulates BMP4, whereas the trophectoderm differentially upregulates WNT3A, consistent with their in-situ expression (Fig 5B, [11]).

**Fig 5:**
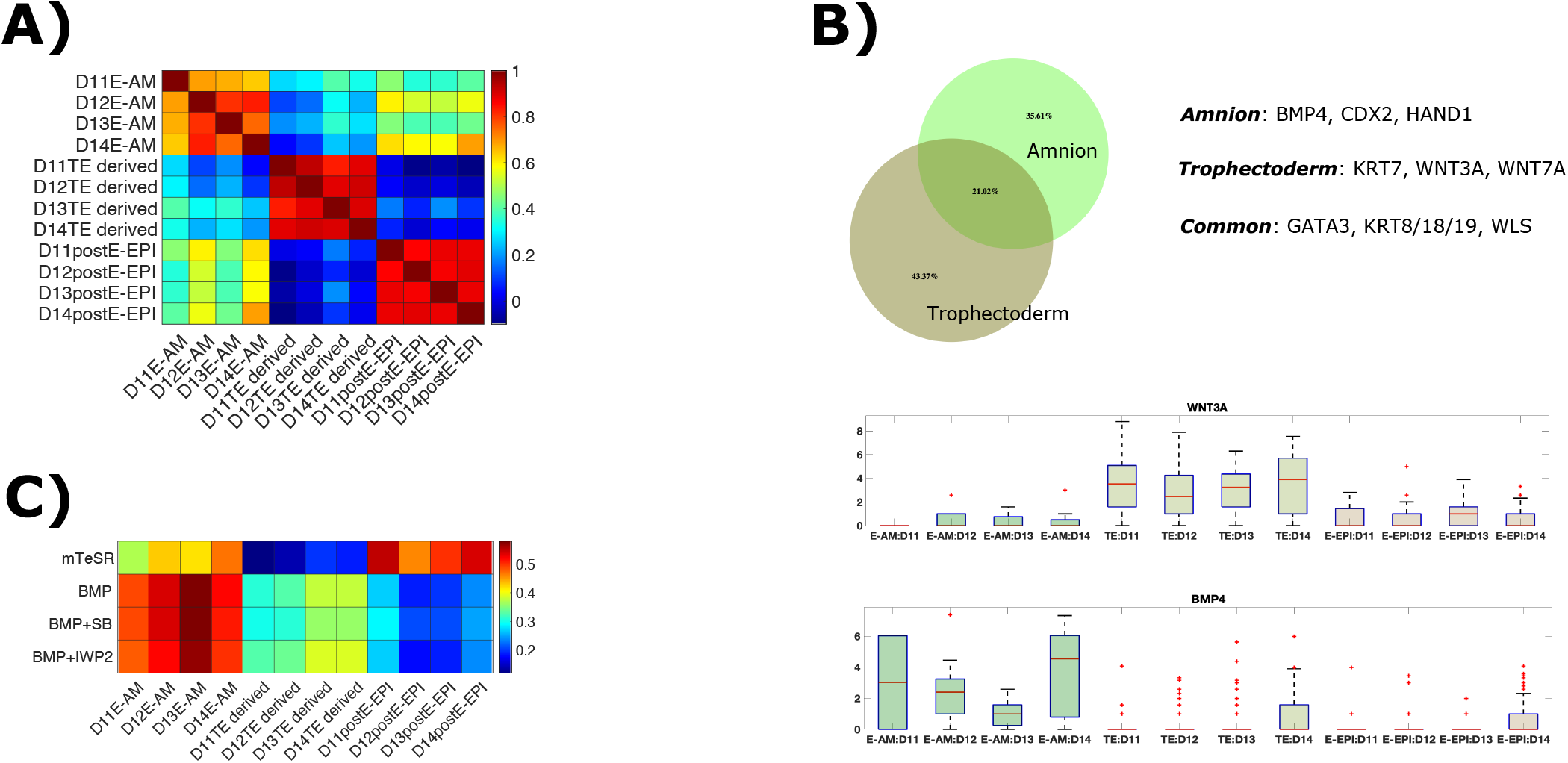
BMP-treated hESCs transcriptionally resemble early post-implantation amnion cells. ***(A)*** Pearson correlation coefficients between indicated samples for 571 lineage-specific genes determined from in vivo monkey embryo data. ***(B)*** Venn diagram for differentially upregulated genes in indicated samples compared to the epiblast. Amnion refers to samples labeled as E-AM, trophectoderm to TE derived and epiblast to postE-epiblast in (A). ***(C)*** Pearson correlation coefficients between indicated samples for 560 lineage-specific genes determined from in vivo monkey embryo data.

Finally, comparing the expression of lineage-specific genes in BMP-treated hESCs and monkey embryo revealed that BMP-treated hESCs are transcriptionally more similar to monkey amnion than monkey trophectoderm-derived cells (Fig 5C). This is intriguing because previous studies have shown that BMP-treated hESCs’ can differentiate towards mature trophectoderm subtypes [3]. Assuming transcriptional similarity between human and monkey amnion, this result suggests that hESCs may possess a remarkable ability to differentiate into mature trophectoderm cells via an amnion-like transcriptional state.

### Xiang et al dataset contains additional mislabeled cellular populations

Correlation analyses of human lineage specific genes across different cell types in the Xiang et al dataset revealed that two additional cell populations – putative intermediate cells and inner cell mass (ICM) cells, are likely mislabeled (Fig S1E, F).

#### Putative intermediate cells are mislabeled extra embryonic mesoderm cells

Putative intermediate cells are a novel cell type identified in the Xiang et al study. In the tSNE analyses presented in Figure 2B in Xiang et al, D12 intermediate cells are placed with epiblast and amnion cell types whereas most D14 intermediate cells are placed in the PSA cluster with amnion and primitive streak cell types. This indicates that intermediate cells represent an epiblast-derived cell population.

However, in the lineage-specific gene space in our analyses, intermediate cells exhibit maximum transcriptional similarity with primitive endoderm cells (Fig S1E, F), not epiblast cells. This trend is preserved at the level of single cells, where a majority of intermediate cells are not transcriptionally correlated with the epiblast or epiblast-derived primitive streak cells (Fig 6A). Consistent with this, most of the intermediate cells do not express pluripotency and primitive streak markers, thus questioning their placement in the PSA cluster (Fig S3A, B).

**Fig 6:**
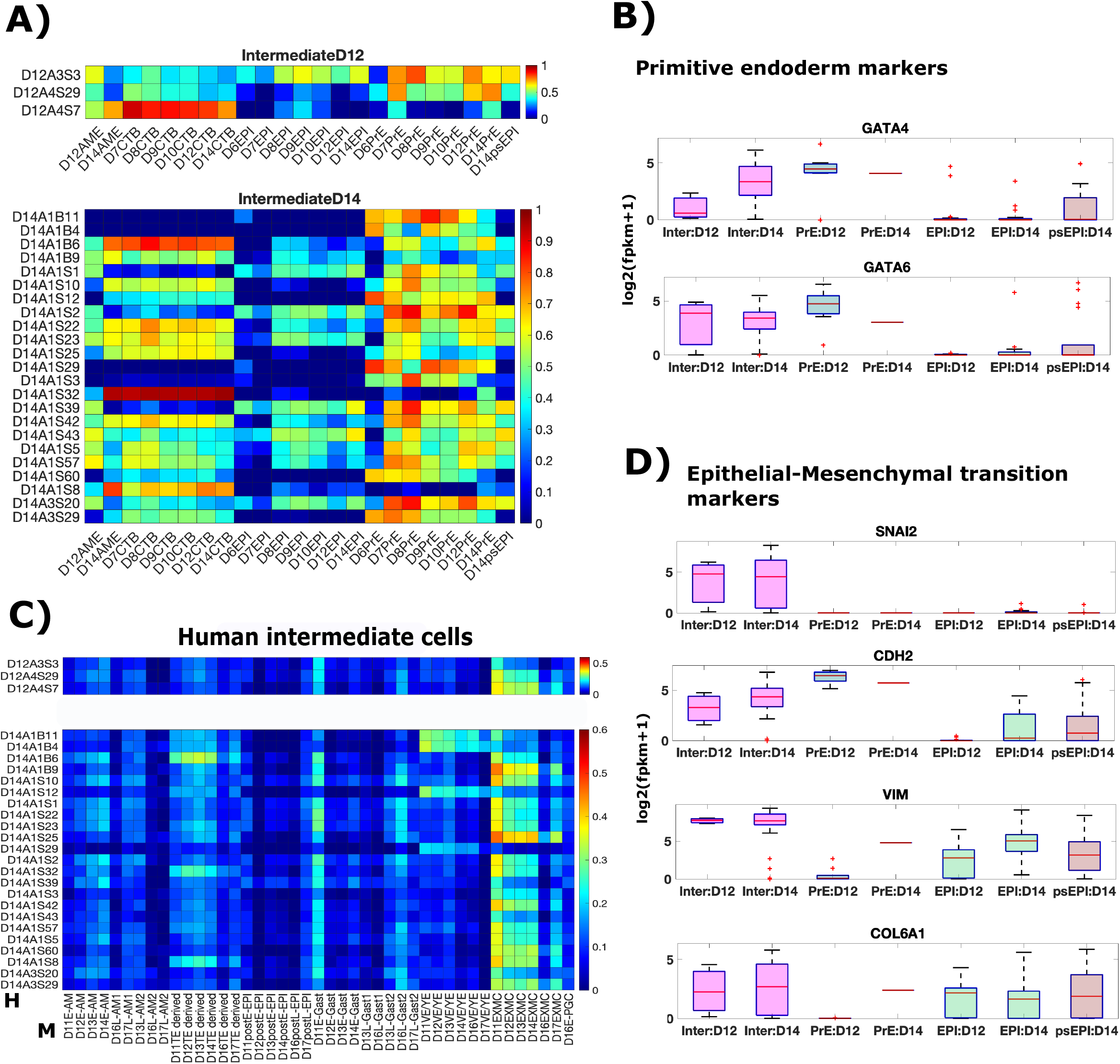
Putative human intermediate cells are mislabeled extra embryonic mesoderm cells. ***(A, C)*** Heatmap showing Pearson correlation coefficients for expression of known lineage markers in human embryo (A) or variable genes in the monkey embryo (C) between individual intermediate cells and indicated cell type averages. Genes in (C) are same as those in Fig S2D ***(B, D)*** Box plots showing expression of indicated genes in indicated lineages.

Most intermediate cells (20/26; 2/3 D12, 18/23 D14) are transcriptionally correlated with primitive endoderm cells (Fig 6A). Consistent with this, intermediate cells express primitive endoderm markers like GATA4/6 at a level comparable to primitive endoderm cells on the same day (Fig 6B). This suggests that intermediate cells represent mislabeled primitive endoderm cells or primitive endoderm derived cells.

Similar to putative amnion cells, the misclassification of putative intermediate cells is also due to the inclusion of pseudogenes. When the principal component analysis is limited to protein coding genes, intermediate cells share PC1 with primitive endoderm cells on both D12 and D14 in all the three gene sets analyzed (Fig 3A, C, D), indicating a high transcriptional similarity of protein coding genes between intermediate cells and primitive endoderm cells.

In the variable gene space of the monkey embryo that separates the two primitive endoderm derived lineages – the EXMC and the VE/YE, most intermediate cells (22/26; 3/3 D12, 19/23 D14) exhibit maximum transcriptional similarity with monkey EXMC cells. Consistent with this, human intermediate cells express known monkey EXMC genes like GATA4, GATA6, COL6A1, VIM, CDH2, SNAI2 (Fig 6B,D, [25] – Extended Figure 5d). It is worth noting that the images in Xiang et. al 2019 do not show a distinct EXMC cell population. However, it is plausible that some primitive endoderm cells have started differentiating towards the EXMC, but a separate EXMC tissue is not yet formed.

#### Putative ICM cells are mislabeled CTB cells

During implantation, the epiblast transitions between the naïve and primed pluripotent states, in both mouse and monkey [8,9]. At the molecular level, this transition results in a reduced expression of naïve pluripotency genes, along with a sustained expression of core pluripotency genes [25,29]. As precursors of epiblast cells, ICM cells are expected to be transcriptionally similar to epiblast cells and express either higher or comparable levels of naïve pluripotency markers as pre-implantation epiblast cells. However, the ICM cells identified in Xiang et al 2019 do not satisfy these conditions.

Comparing expression of lineage specific genes in individual putative ICM cells with average expression of those genes in different lineages in the embryo reveals that a majority of these cells (49/52) are transcriptionally correlated with CTB, not epiblast cells (Fig 7A). Consistent with this, putative ICM cells express other known trophoblast markers – TP63, TFAP2A, CDX2 at a level comparable with CTB cells on the same days (Fig 7B). On the other hand, these cells do not express core pluripotency markers – NANOG, SOX2 and OCT4 (Fig 7C). This data contradicts previous literature which shows that D6-9 ICM cells express pluripotency, not trophectoderm genes [18–21,23].

**Fig 7:**
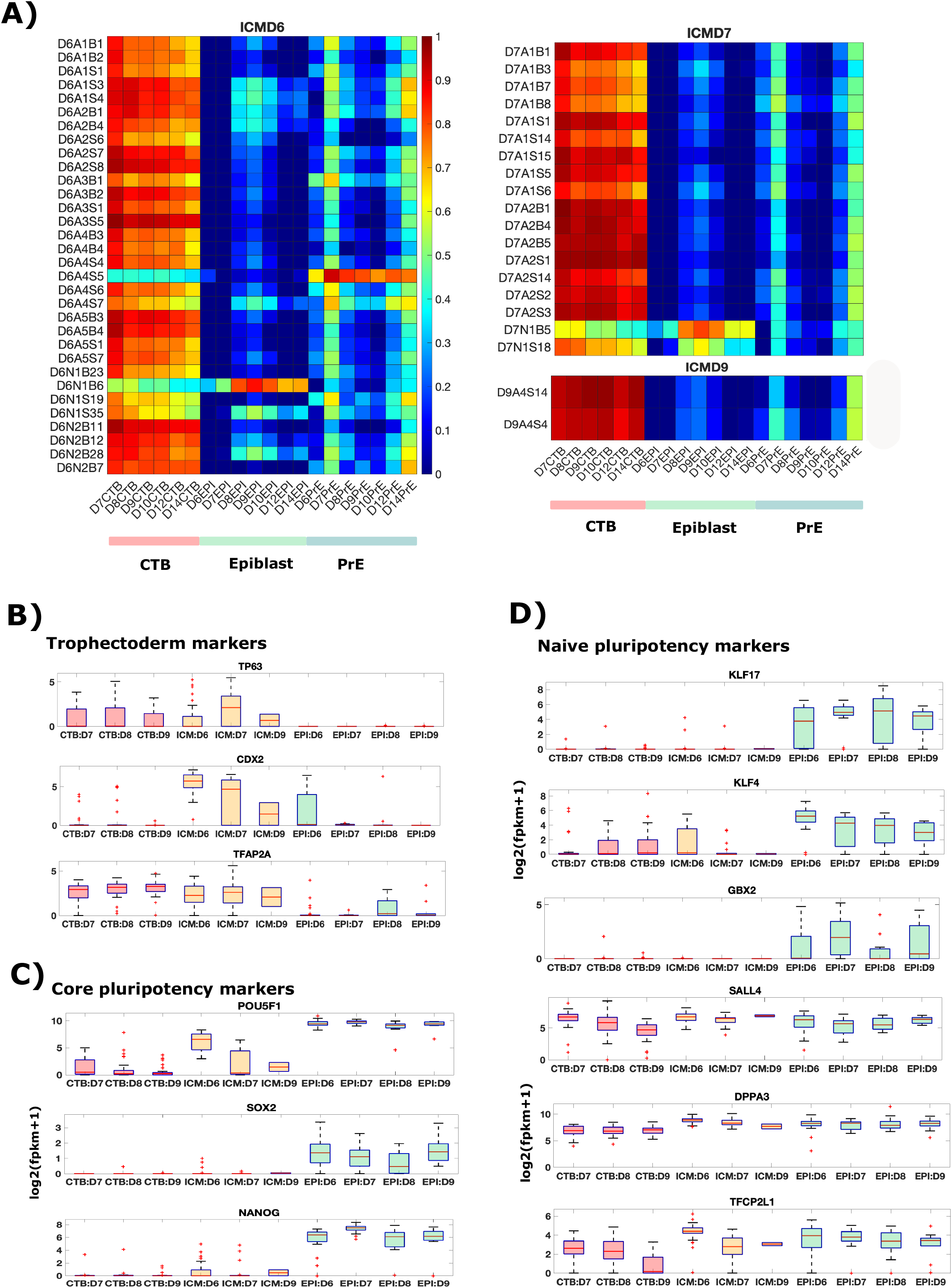
Putative human ICM cells are mislabeled CTB cells. ***(A)*** Heatmap showing Pearson correlation coefficients for expression of known lineage markers between individual ICM cells and indicated cell type averages. ***(B, C, D)*** Box plots showing expression of indicated genes in indicated lineages.

Xiang et al 2019 claim that they observe a gradual maturation of the epiblast from the naïve to the primed pluripotency state (Xiang et al 2019, Extended Figure 9). However, naïve pluripotency markers – KLF17, KLF4, GBX2 are expressed in fewer putative ICM cells and at a lower level, compared with epiblast cells. Other naïve pluripotency markers, SALL4 and DPPA3 are expressed in comparable levels in ICM, epiblast and CTB cells, indicating that these markers are not specific to ICM/epiblast. The only exception is TFCP2L1, which exhibits a slightly higher median expression in day 6 putative ICM cells, compared to epiblast and CTB cells (Fig 7D). However, TFCP2L1 protein is expressed at comparable levels in both CTB and ICM cells of the day 6 human embryos (Xiang et al 2019, Extended data Figure 9a), indicating that TFCP2L1 is also not specific to ICM/epiblast. Taken together, this data shows that the cells labelled as ICM express naïve pluripotency markers at a level comparable to CTB cells, and lower than epiblast cells.

In D6-9 human embryos, the absolute number of trophectoderm cells is higher than ICM/epiblast cells [18–21,23]. Thus, it is surprising to obtain 32 ICM cells, 28 epiblast cells and 0 CTB in D6 human embryos (Fig S1B). It is more likely that 30/32 ICM cells, which exhibit transcriptional similarity with CTB cells, are mislabeled CTB cells (Fig 7A).

It is worth noting that the cells of the D5 human embryo cannot be distinguished based on the known lineage markers as they co-express markers of the three lineages ([17]). However, most of the putative ICM cells in this dataset (49/52) clearly correlate more with trophoblast cells than the other two lineages and express trophoblast markers on a par with CTB cells, indicating that they do not correspond to the early heterogenous population and are likely mislabeled CTB cells.

Taken together, the above analyses suggest that 49/52 ICM cells – 30/32 Day6, 17/18 Day7 and 2/2 Day9 ICM cells are mislabeled CTB cells on the corresponding days. Of the remaining 3 cells, 2 represent epiblast cells and 1 represents primitive endoderm on the corresponding day, as indicated by the transcriptional similarity of known lineage genes (Fig 7A).

## Discussion

Human embryonic stem cells (hESCs) offer a unique opportunity to probe early stages of human development. Their immense potential to differentiate into a variety of different cell types offers a valuable resource for both translational and fundamental research. However, to make accurate inferences, it is essential to determine the identity of cells obtained by differentiation of hESCs through comparisons with embryos in vivo.

The identity of CDX2 positive cells obtained after treating hESCs with BMP4 for 2-4 days, has remained controversial. They have been identified as trophectoderm, extra-embryonic mesoderm and amnion cells ([3–5]). Previously, we showed that these cells are transcriptionally similar to human trophectoderm, consistent with their expression of known trophectoderm markers and ability to differentiate towards mature trophectoderm subtypes[3,14]. Additionally, we argued that these cells cannot represent extra-embryonic mesoderm fate as they do not express GATA4/6, key markers associated with monkey extra-embryonic mesoderm [6,25]. The lack of in vivo amnion data and the expression of known trophectoderm markers in the putative in vitro amnion cells [4], has prevented a direct comparison with in vivo amnion.

In this study, we carefully examined the recently published putative human amnion data [10] to discern the unique transcriptional signature of amnion cells. Our analyses revealed that the inclusion of pseudogenes leads to the mislabeling of putative amnion cells. In the absence of pseudogenes, amnion cells do not form a separate cluster in the principal component analyses and are instead spread across the principal component space, with most of the cells in the trophoblast region (Fig 3C). Restricting the analyses to amnion and trophectoderm cell types further revealed that most of the amnion cells transcriptionally resemble syncytiotrophoblast cells (Fig 4). The erroneous results obtained when pseudogene expression is not excluded highlight the need to carefully compile appropriate lists of genes to compare different cell populations. A general method for doing so is an important topic for future study.

In the absence of a unique human amnion transcriptome, we utilized the early post-implantation monkey embryo to determine the similarity between BMP-treated hESCs and amnion cells. Our analyses revealed that BMP-treated hESCs are transcriptionally more similar to the amnion than trophectoderm cells in the monkey embryo (Fig 5). Together with previous results [3,6,14], this result indicates that hESCs may possess a remarkable ability to differentiate towards a mature trophectoderm state via an amnion-like state, and presumably the transcriptional similarity between amnion and trophectoderm cells enables this route of differentiation. To verify if this ability extends to cells in an intact human or monkey embryo, one could transplant amnion cells in the trophectoderm region and examine if they continue to develop as trophectodermal cells.

We also found additional mislabeled cellular populations in the Xiang et al dataset. One of these is the intermediate cell population. Although Xiang et al do not comment on their in vivo identity, their placement with the epiblast and amnion cells implies that they presumably represent an epiblast derived cell population. However, we show that excluding pseudogenes changes their position in the principal component space and moves them closer to the primitive endoderm cluster, indicating that they likely represent primitive-endoderm derived cells (Fig 3C). Comparison with monkey embryo revealed that they likely represent extra-embryonic mesoderm cells which are known to express key primitive endoderm genes (Fig 6C, [25]). It is worth noting that there is no morphological extra-embryonic region covering the amnionembryo-primitive endoderm region in the Xiang et al study [10]. However, it is plausible that some cells have started differentiating towards the extra-embryonic mesoderm but have not occupied their morphological location yet. In the future, time lapse imaging studies could discern the precise dynamics of extra-embryonic mesoderm specification and migration.

## Methods

In this analyses, we utilized the transcriptome data of the human and monkey embryo published previously [10,13]. All the analyses were performed at the level of genes. For genes with multiple transcripts, cumulative expression of all transcripts was considered as the gene read count and the ensembl gene id of most expressed transcript was considered as its gene id. PCA and correlation analyses within a given dataset was performed on log transformed read counts (log2 (FPKM+1)). For all analyses except in Fig 5, genes were selected based on expression counts (FPKM>1 in at least 50% cells of a given lineage) and variability across cells (CV threshold). To determine lineage specific genes in Fig 8, we used EBSeq with an FDR cutoff of 0.01 for pairwise differential gene analyses in order to reduce the overlap of lineage-specific genes across lineages [30]. CV thresholds and number of genes are indicated in relevant figures and figure legends. For analyses between the two datasets, each dataset was filtered to exclude non-expressed genes (FPKM = 0 in all cells within a dataset), after which the log normalized read counts (log2(FPKM+1)) were transformed into z scores. Only genes retained in both datasets after filtering were utilized for correlation analyses. Human gene orthologs of monkey genes were obtained from Supplementary information in [25].

---

**Fig S1:**
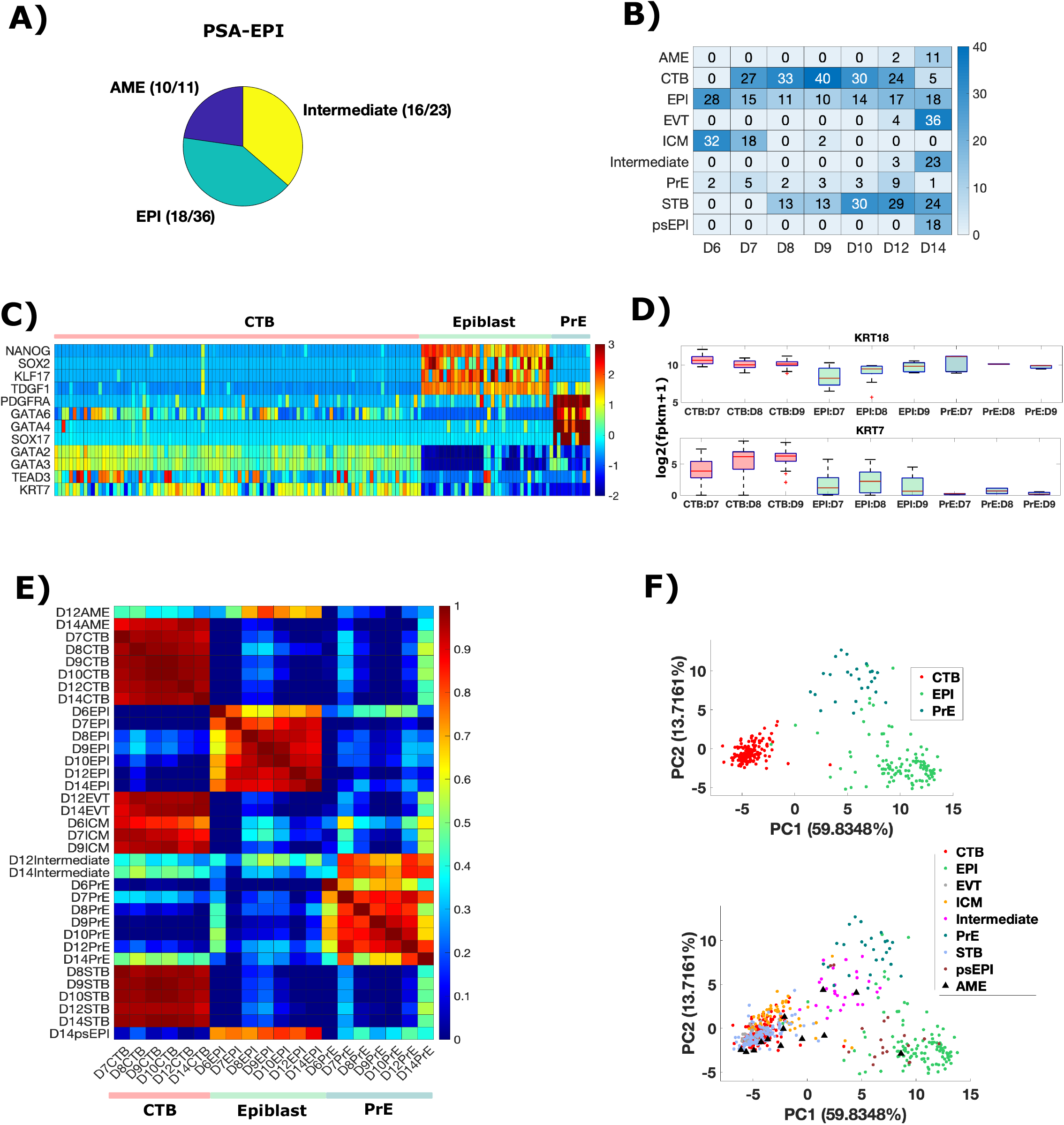
Known lineage markers separate human epiblast, trophectoderm and primitive endoderm lineages. ***(A)*** Distribution of cells in primitive streak anlage cluster. ***(B)*** No. of cells corresponding to each cell type on each day. psEPI cells are epiblast cells in the primitive streak anlage cluster. Data in A and B is based on information in Supplementary table 8 (S8.1, S8.3) in Xiang et al study. ***(C)*** Heatmap showing expression of known lineage markers in each cell in D7-9 cytotrophoblast (CTB), epiblast (EPI) and primitive endoderm (PrE) lineages. Values correspond to z-scores of indicated genes. Z-scores were calculated for each gene across all cells in D7-9 EPI, PrE and CTB lineages. ***(D)*** Box plots showing expression of indicated genes in indicated lineages. ***(E)*** Heatmap showing Pearson correlation coefficients of average expression of known lineage markers in each cell type. ***(F)*** PCA of known lineage marker gene expression across all 555 cells. In the top plot, a subset of cells corresponding to CTB, EPI and PrE lineages are shown. The number of genes used in analyses is indicated above the plot.

**Fig S2:**
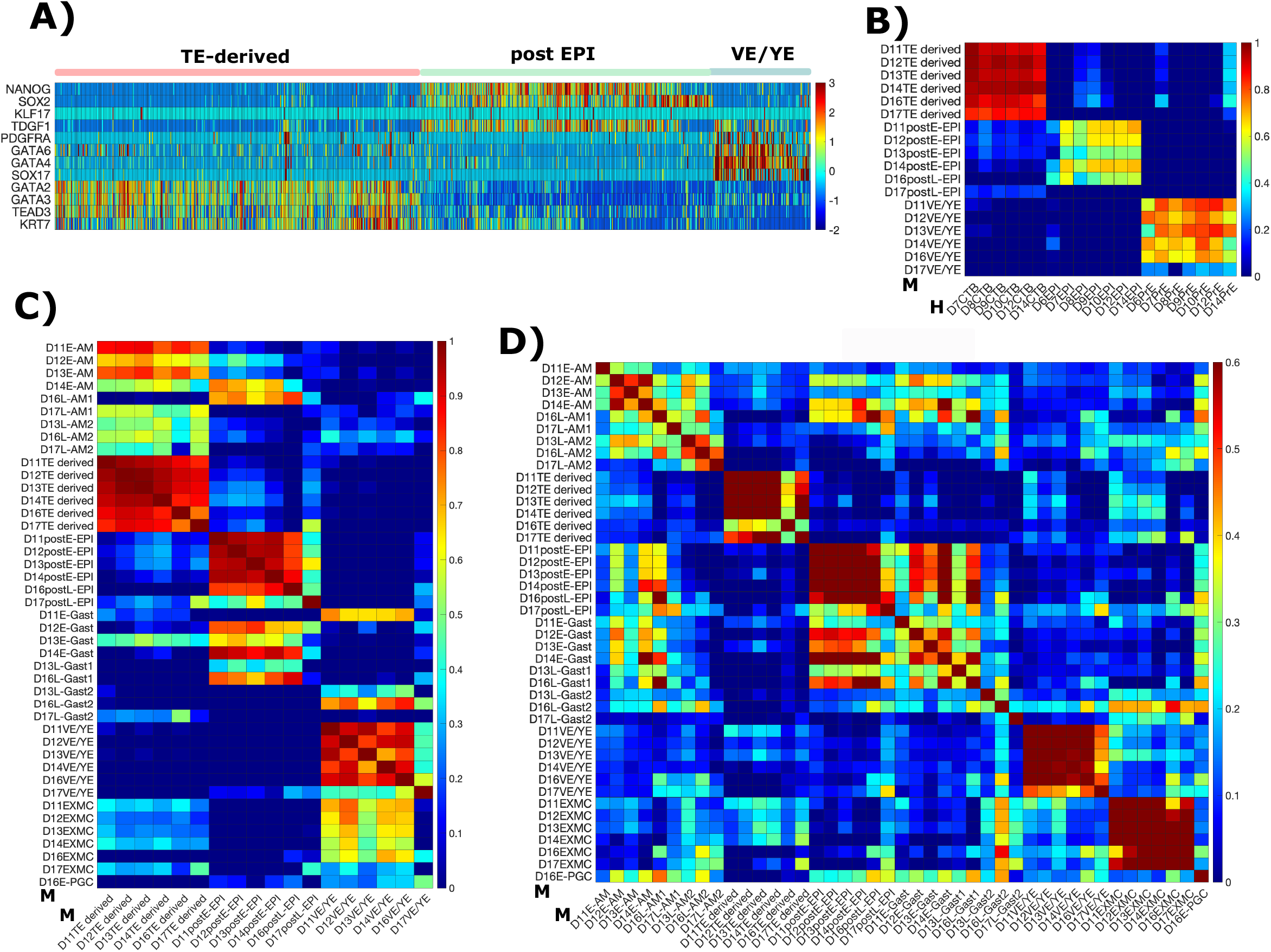
Human lineage specific genes delineate three primary lineages of the monkey embryo but do not distinguish amnion and trophectoderm lineages. ***(A)*** Heatmap showing expression of human known lineage markers (same genes as in Fig1C) in each cell in D11-D17 trophectoderm-derived, post implantation epiblast and visceral endoderm/yolk-sac endoderm (VE/YE) lineages in the monkey embryo. Values correspond to z-scores of indicated genes. Z-scores were calculated for each gene across all cells in the plotted lineages. ***(B-D)*** Heatmap showing Pearson correlation coefficients of average expression of either known lineage markers (B, C) or variable genes in the monkey embryo ((D); CV>1 across 1453 monkey cells; 2440 genes) in indicated cell types. The symbols H and M represent the parent embryo of cells - human(H), monkey(M).

**Fig S3:**
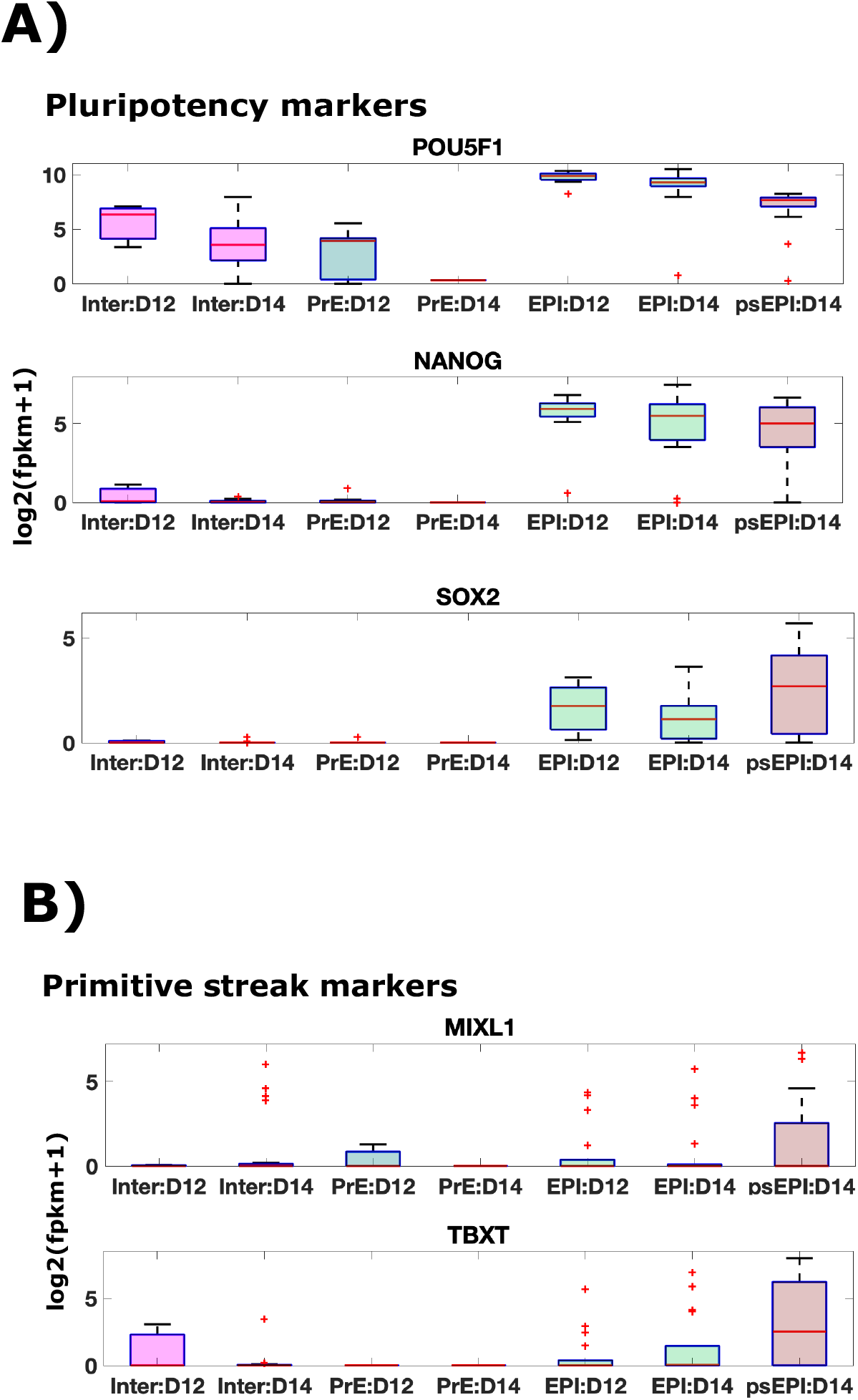
Intermediate cells do not express pluripotency and primitive streak markers. ***(A, B))*** Box plots showing expression of indicated genes in indicated lineages.

**Supplementary Table 1** Gene lists used for analyses in the indicated Figures.

## Notes

### Competing Interest Statement

The authors have declared no competing interest.

## References

1. Dupont G, Yilmaz E, Loukas M, Macchi V, De Caro R, Tubbs RS. Human embryonic stem cells: Distinct molecular personalities and applications in regenerative medicine. Clin Anat. 2019. 32(3):354–60. doi:10.1002/ca.23318

2. Fu J, Warmflash A, Lutolf MP. Stem-cell-based embryo models for fundamental research and translation. Nature Materials. Nature Research; 2020. doi:10.1038/s41563-020-00829-9

3. Xu R-H, Chen X, Li DS, Li R, Addicks GC, Glennon C, et al. BMP4 initiates human embryonic stem cell differentiation to trophoblast. Nat Biotechnol. 2002. 20(12):1261–4. doi:10.1038/nbt761

4. Shao Y, Taniguchi K, Gurdziel K, Townshend RF, Xue X, Yong KMA, et al. Selforganized amniogenesis by human pluripotent stem cells in a biomimetic implantation-like niche. Vol. 16, Nature Materials. Nature Publishing Group; 2017. p. 419–27. doi:10.1038/NMAT4829

5. Bernardo AS, Faial T, Gardner L, Niakan KK, Ortmann D, Senner CE, et al. BRACHYURY and CDX2 Mediate BMP-Induced Differentiation of Human and Mouse Pluripotent Stem Cells into Embryonic and Extraembryonic Lineages. Cell Stem Cell. 2011. 9(2):144–55. doi:10.1016/j.stem.2011.06.015

6. Chhabra S, Liu L, Goh R, Kong X, Warmflash A. Dissecting the dynamics of signaling events in the BMP, WNT, and NODAL cascade during self-organized fate patterning in human gastruloids. Lowell S, editor. PLOS Biol. 2019. 17(10):e3000498. doi:10.1371/journal.pbio.3000498

7. Rossant J, Boroviak T, Loos R, Bertone P, Smith A, Nichols J, et al. Mouse and human blastocyst-derived stem cells: vive les differences. Development. 2015. 142(1):9–12. doi:10.1242/dev.115451

8. Dobreva MP, Pereira PNG, Deprest J, Zwijsen A. On the origin of amniotic stem cells: of mice and men. Int J Dev Biol. 2010. 54(5):761–77. doi:10.1387/ijdb.092935md

9. Kinder SJ, Tsang TE, Quinlan GA, Hadjantonakis AK, Nagy A, Tam PP. The orderly allocation of mesodermal cells to the extraembryonic structures and the anteroposterior axis during gastrulation of the mouse embryo. Development. 1999.126(21):4691–701.

10. Xiang L, Yin Y, Zheng Y, Ma Y, Li Y, Zhao Z, et al. A developmental landscape of 3D-cultured human pre-gastrulation embryos. Nature. 2019. 577(7791):537–42. doi:10.1038/s41586-019-1875-y

11. Sasaki K, Nakamura T, Okamoto I, Yabuta Y, Iwatani C, Tsuchiya H, et al. The Germ Cell Fate of Cynomolgus Monkeys Is Specified in the Nascent Amnion. Dev Cell. 2016. 39(2):169–85. doi:10.1016/j.devcel.2016.09.007

12. Knöfler M, Haider S, Saleh L, Pollheimer J, Gamage TKJB, James J. Human placenta and trophoblast development: key molecular mechanisms and model systems. Vol. 76, Cellular and Molecular Life Sciences. Birkhauser Verlag AG; 2019. p. 3479–96. doi:10.1007/s00018-019-03104-6

13. Ma H, Zhai J, Wan H, Jiang X, Wang X, Wang L, et al. In vitro culture of cynomolgus monkey embryos beyond early gastrulation. Science (80-). 2019. 366(6467). doi:10.1126/science.aax7890

14. Li Y, Moretto-Zita M, Soncin F, Wakeland A, Wolfe L, Leon-Garcia S, et al. BMP4-directed trophoblast differentiation of human embryonic stem cells is mediated through a ΔNp63+ cytotrophoblast stem cell state. Dev. 2013. 140(19):3965–76. doi:10.1242/dev.092155

15. Luckett WP. Origin and differentiation of the yolk sac and extraembryonic mesoderm in presomite human and rhesus monkey embryos. Am J Anat. 1978. 152(1):59–97. doi:10.1002/aja.1001520106

16. Enders AC, King BF. Formation and differentiation of extraembryonic mesoderm in the rhesus monkey. Am J Anat. 1988. 181(4):327–40. doi:10.1002/aja.1001810402

17. Stirparo GG, Boroviak T, Guo G, Nichols J, Smith A, Bertone P. Integrated analysis of single-cell embryo data yields a unified transcriptome signature for the human preimplantation epiblast. Dev. 2018. 145(3). doi:10.1242/dev.158501

18. Roode M, Blair K, Snell P, Elder K, Marchant S, Smith A, et al. Human hypoblast formation is not dependent on FGF signalling. Dev Biol. 2012. 361(2):358–63. doi:10.1016/j.ydbio.2011.10.030

19. Niakan KK, Eggan K. Analysis of human embryos from zygote to blastocyst reveals distinct gene expression patterns relative to the mouse. Dev Biol. 2013. 375(1):54–64. doi:10.1016/j.ydbio.2012.12.008

20. Deglincerti A, Croft GF, Pietila LN, Zernicka-Goetz M, Siggia ED, Brivanlou AH. Self-organization of the in vitro attached human embryo. Nature. 2016. 533(7602):251–4. doi:10.1038/nature17948

21. Shahbazi MN, Jedrusik A, Vuoristo S, Recher G, Hupalowska A, Bolton V, et al. Self-organization of the human embryo in the absence of maternal tissues. Nat Cell Biol. 2016. 18(6):700–8. doi:10.1038/ncb3347

22. Chen AE, Egli D, Niakan K, Deng J, Akutsu H, Yamaki M, et al. Optimal Timing of Inner Cell Mass Isolation Increases the Efficiency of Human Embryonic Stem Cell Derivation and Allows Generation of Sibling Cell Lines. Vol. 4, Cell Stem Cell. Cell Stem Cell; 2009. p. 103–6. doi:10.1016/j.stem.2008.12.001

23. Blakeley P, Fogarty NME, del Valle I, Wamaitha SE, Hu TX, Elder K, et al. Defining the three cell lineages of the human blastocyst by single-cell RNA-seq. Development. 2015. 142(18):3151–65. doi:10.1242/dev.123547

24. Petropoulos S, Edsgärd D, Reinius B, Deng Q, Panula SP, Codeluppi S, et al. Single-Cell RNA-Seq Reveals Lineage and X Chromosome Dynamics in Human Preimplantation Embryos. Cell. 2016. 165(4):1012–26. doi:10.1016/j.cell.2016.03.023

25. Nakamura T, Okamoto I, Sasaki K, Yabuta Y, Iwatani C, Tsuchiya H, et al. A developmental coordinate of pluripotency among mice, monkeys and humans. Nature. 2016. 537(7618):57–62. doi:10.1038/nature19096

26. Pink RC, Wicks K, Caley DP, Punch EK, Jacobs L, Carter DRF. Pseudogenes: Pseudo-functional or key regulators in health and disease. Vol. 17, RNA. Cold Spring Harbor Laboratory Press; 2011. p. 792–8. doi:10.1261/rna.2658311

27. Milligan MJ, Lipovich L. Pseudogene-derived lncRNAs: emerging regulators of gene expression. Front Genet. 2015.5(FEB):476. doi:10.3389/fgene.2014.00476

28. Okae H, Toh H, Sato T, Hiura H, Takahashi S, Shirane K, et al. Derivation of Human Trophoblast Stem Cells. Cell Stem Cell. 2018. 22(1):50–63.e6. doi:10.1016/j.stem.2017.11.004

29. Nichols J, Smith A. Naive and Primed Pluripotent States. Vol. 4, Cell Stem Cell. Cell Stem Cell; 2009. p. 487–92. doi:10.1016/j.stem.2009.05.015

30. Leng N, Dawson JA, Thomson JA, Ruotti V, Rissman AI, Smits BMG, et al. EBSeq: an empirical Bayes hierarchical model for inference in RNA-seq experiments. Bioinformatics. 2013. 29(8):1035–43. doi:10.1093/bioinformatics/btt087

